# The Landscape of *Parkin* Variants Reveals Pathogenic Mechanisms and Therapeutic Targets in Parkinson’s Disease

**DOI:** 10.1101/445551

**Authors:** Wei Yi, Emma J. MacDougall, Matthew Y. Tang, Andrea I. Krahn, Ziv Gan-Or, Jean-François Trempe, Edward A. Fon

## Abstract

Mutations in *Parkin* (*PARK2*), which encodes an E3 ubiquitin ligase implicated in mitophagy, are the most common cause of early onset Parkinson’s Disease (PD). Hundreds of naturally occurring *Parkin* variants have been reported, both in PD patient and population databases. However, the effects of the majority of these variants on the function of Parkin and in PD pathogenesis remains unknown. Here we develop a framework for classification of the pathogenicity of *Parkin* variants based on the integration of clinical and functional evidence – including measures of mitophagy and protein stability, and predictive structural modeling – and assess 51 naturally occurring *Parkin* variants accordingly. Surprisingly, only a minority of *Parkin* variants, even among those previously associated with PD, disrupted Parkin function. Moreover, a few of these naturally occurring *Parkin* variants actually enhanced mitophagy. Interestingly, impaired mitophagy in several of the most common *pathogenic* Parkin variants could be rescued both by naturally-occurring (p.V224A) and structure-guided designer (p.W403A; p.F146A) hyperactive Parkin variants. Together, the findings provide a coherent framework to classify *Parkin* variants based on pathogenicity and suggest that several *pathogenic Parkin* variants represent promising targets to stratify patients for genotype-specific drug design.

## Introduction

Parkinson’s disease (PD) is the second most common neurodegenerative disease. Although most PD cases are sporadic, a fraction are familial and caused by mutations in different genes (1). Mutations in the *Parkin* (*PARK2*) gene are the most common cause of autosomal recessive early-onset parkinsonism (EOPD) and are believed to result in a loss of Parkin protein function (2). *Parkin* variants include rearrangements and copy number variations, such as deletions and duplications of exons, as well as single nucleotide variants (SNVs) that cause missense, nonsense, or splice site mutations (3-5). Of these, missense variants are the most frequently reported in PD patients and, because they likely impede Parkin protein function rather than disrupting protein expression, may represent viable targets for therapies that enhance Parkin activity.

To envisage such genotype-specific therapies, the pathogenicity of the many *Parkin* variants in the population first needs to be determined. The American College of Medical Genetics and Genomics for Molecular Pathology (ACMG-AMP) has outlined five standard terminologies to describe variants identified in genes that cause Mendelian disorders (6). “*Pathogenic*” and “*likely pathogenic*” indicate a clear or very likely disease-causing effect of a variant, respectively. Conversely,“*likely benign*,” and “*benign*” indicate variants that are not disease-causing. Variants that cannot be assigned to one of these four groups are designated as “*uncertain significance*”. Over 200 *Parkin* missense variants have been deposited in public repositories (4, 5, 7, 8). However, to date, only a minority have been clearly annotated based on formal criteria.

A clear assignment of a *Parkin* variant requires integrating different lines of evidence that fall into two broad categories (9). Clinical evidence consists of the association or segregation of the variant with disease (or the absence of) in human cohorts or within families with multiple affected individuals. Functional evidence refers to the consequence(s) of the variant, using experimental assays that measure biochemical and cellular properties, as well as computational algorithms that model the effect(s) of the variant based on protein structure and function. Clinical evidence has a hierarchical relationship relative to functional evidence and prevails when a discrepancy or conflict arises between clinical and functional observations (9, 10).

Parkin is a basally autoinhibited E3 ubiquitin (Ub) ligase, which contains an N-terminal Ub-like (Ubl) domain, connected through a linker to four zinc-coordinating domains, RING0, RING1, In-Between-RING (IBR), and RING2, which form a core designated as the R0RBR (11). Parkin is activated by PINK1, a mitochondrial kinase that is also implicated in EOPD (1). PINK1 accumulates on damaged mitochondria upon depolarization, where it phosphorylates nearby Ub (12-15). Parkin binds to phospho-Ub (pUb), which recruits Parkin to mitochondria and facilitates PINK1 phosphorylation of the Parkin Ubl, which in turn fully activates Parkin (16-18). Parkin then ubiquitinates multiple outer mitochondrial membrane targets, triggering a feed-forward amplification loop, that leads to the clearance of damaged mitochondria via autophagy (mitophagy) (reviewed in (19)). Some *Parkin* missense variants found in patients have been shown to affect protein folding, Parkin autoubiquitination, protein-protein interactions or recruitment to mitochondria (20-25). However, as the pathogenic nature of most of these variants was not clear, the disease relevance of the functional alteration remained to be determined.

Here, we characterized all *Parkin* missense variants found in public databases, according to ACMG-AMP criteria. We then used a cell-based assay to quantify Parkin-mediated mitophagy and Parkin protein levels. We also applied structural simulations to explore the mechanisms underlying the observed functional alterations. Integrating these data, we find that only a minority of *Parkin* variants can be considered *pathogenic*. Interestingly, we identified several naturally-occurring *Parkin* variants in the population that increase mitophagy in cells. Remarkably, such hyperactive *Parkin* variants were able to rescue the impaired function of several common *pathogenic Parkin* variants. Our study suggests that several *pathogenic* PD-linked *Parkin* mutations represent promising targets amenable to genotype-specific drug design.

## Results

### Most *Parkin* missense variants lack sufficient clinical evidence to establish pathogenicity

To classify *Parkin* variants, we utilized Sherloc (semiquantitative, hierarchical evidence-based rules for locus interpretation), a classification framework that translates the ACMG-AMP standards to a set of discrete, but related, rules with refined weights (9). Clinical evidence was examined first, as it most directly relates to disease (9). Briefly, data from population databases, including minor allele frequency (MAF) and homozygote counts, and clinical records in PD-specific databases that report PD patients or unaffected family members carrying missense variants in *Parkin*, were examined to assign weighted points as pathogenic or benign to each variant (Supplementary Fig. 1). The pathogenic points and benign points were summed-up separately and compared to preset thresholds to assign the variants to one of the five ACMG-AMP categories (Supplementary Fig. 2).

**Figure 1.**
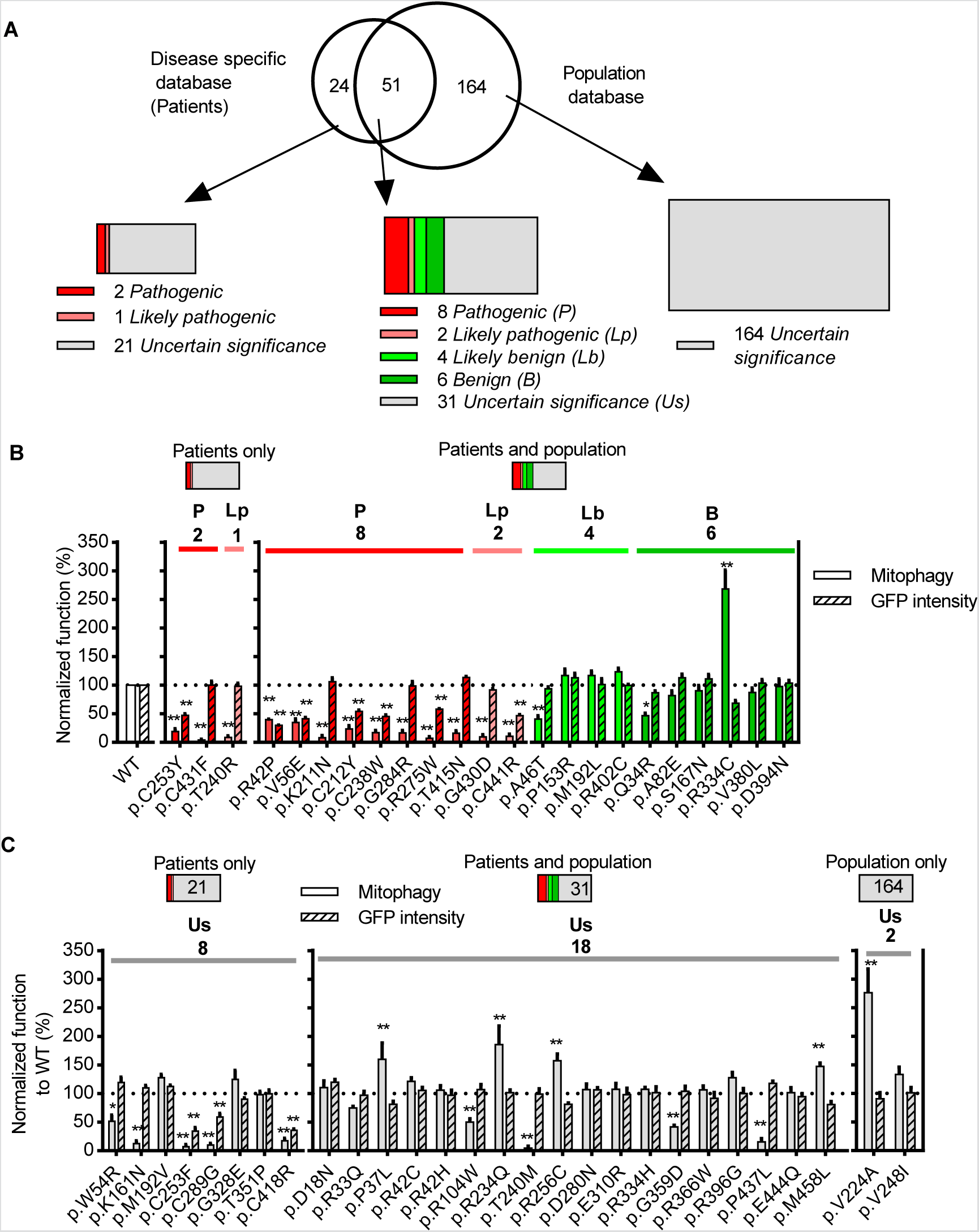
*Parkin* missense variants displayed a wide range of functional alterations. **(A)** 75 *Parkin* missense variants were reported in disease specific databases (PDmutDB, MDSgene), 215 were reported in population databases (ExAC, dbSNP) and 51 were reported in both. Variants were assigned with 1 of 5 standard ACMG terminologies: *Pathogenic* (red), *likely pathogenic* (pink), *likely benign* (green), *benign* (olive) and *uncertain significance* (grey). **(B-C)** Quantification of the function of Parkin missense variants assigned as **(B)** *Pathogenic* (red), *likely pathogenic* (pink), *likely benign* (green), *benign* (olive) or **(C)** Uncertain significance (grey) by clinical evidence. Solid bars show mitophagy after 4h of CCCP treatment quantified from mtKeima signal in U2OS cells expressing GFP-Parkin variants normalized to wildtype (WT) Parkin. Hatched bars show GFP intensity of cells expressing GFP-Parkin variants normalized to WT Parkin. * P<0.05, ** P<0.01, in one-way ANOVA with Dunnett’s post-hoc test comparing the function of each variant with WT. N=3-7.

**Figure 2.**
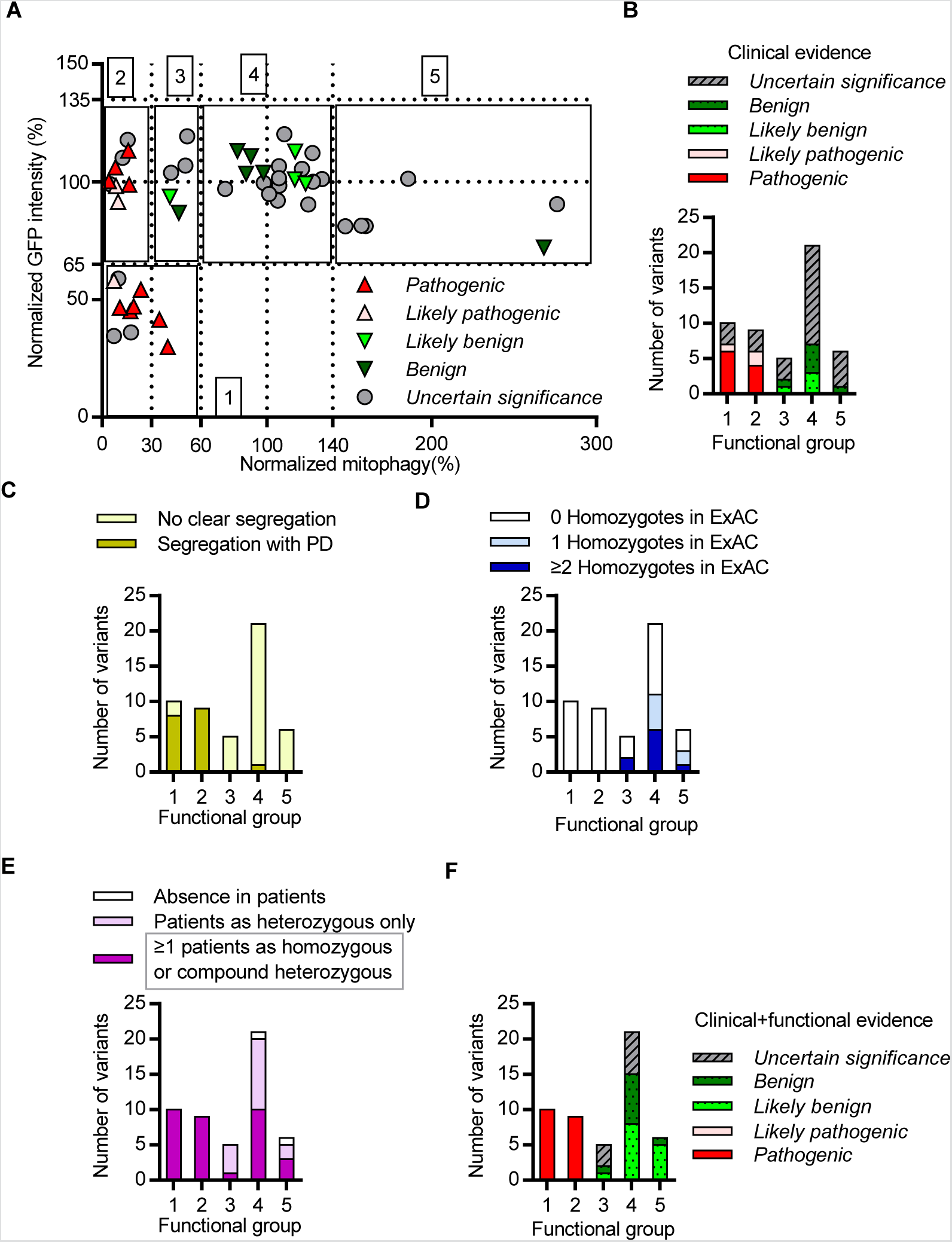
Integration of clinical and functional evidence refined the classification of Parkin variants. **(A)** Functional alteration of variants assigned as *pathogenic* (red), *likely pathogenic* (pink), *likely benign* (green), *benign* (olive), and of *uncertain significance* (grey) based on clinical evidence were plotted for mitophagy activity on the X axis and GFP intensity on the Y axis. Functional alteration segregated into five groups, indicated by black boxes. 1. Significantly decreased mitophagy activity and GFP intensity compared with WT. 2. Severely decreased mitophagy activity with WT GFP intensity. 3. Moderately decreased mitophagy activity with WT GFP intensity. 4. WT mitophagy activity and GFP intensity. 5. Significantly increased mitophagy activity with WT GFP intensity. **(B)** Quantification of the variants within each of the functional groups from (A). **(C)** Quantification of variants within the functional groups described in (A) according to their segregation or lack of segregation with PD in families. **(D)** Quantification of variants within the functional groups described in (A) according to the observation of the variant as more than 1 homozygote (blue), 1 homozygote (light-blue), or no homozygotes (white) in ExAC. **(E)** Quantification of variants within the functional groups described in (A) according to the observation of the variant in PD patients. **(F)** Quantification of variants within the functional groups described in (A) according to their classification based on clinical and functional evidence.

From the PD-specific databases, PDmutDB (3, 4) and MDSgene (5, 26), we identified a total of 75 *Parkin* missense variants in PD patients (Fig. 1A). Most were absent or very rare in the control cohorts from the original reports, possibly due to the small sizes of the control cohorts. Therefore, we searched for the 75 variants in public population databases and found that 51 of them were reported in dbSNPs (7) and in the Exome Aggregation Consortium (ExAC), including high-quality variant calls across 60,706 human exomes (8) (Fig. 1A). In ExAC, we also found an additional 164 *Parkin* missense variants (Fig. 1A). The classification of the missense variants using clinical evidence allowed us to clearly designate thirteen variants in PD as *pathogenic* or *likely pathogenic* and ten variants as *likely benign* or *benign* (Fig. 1A and Table 1). The details of the points assigned to each variant are summarized in Supplementary Table 1. Remarkably, large numbers of remaining variants lacked sufficient data to be assigned to either the *benign* or *pathogenic* categories and were therefore designated as of *uncertain significance* (Fig. 1A, Table 1 and Supplementary Table 1).

**Table 1.** Annotation of *Parkin* missense variants by ACMG terminologies.

| Index | Databases | Parkin Variant | Annotation with clinical evidence | Functional group | Annotation with clinical and functional evidence |
| --- | --- | --- | --- | --- | --- |
| 1 | Disease and population | p.R42P | *Pathogenic* | 1 | *Pathogenic* |
| 2 | Disease and population | p.V56E | *Pathogenic* | 1 | *Pathogenic* |
| 3 | Disease and population | p.K211N | *Pathogenic* | 2 | *Pathogenic* |
| 4 | Disease and population | p.C212Y | *Pathogenic* | 1 | *Pathogenic* |
| 5 | Disease and population | p.C238W | *Pathogenic* | 1 | *Pathogenic* |
| 6 | Disease | p.C253Y | *Pathogenic* | 1 | *Pathogenic* |
| 7 | Disease and population | p.G284R | *Pathogenic* | 2 | *Pathogenic* |
| 8 | Disease and population | p.T415N | *Pathogenic* | 2 | *Pathogenic* |
| 9 | Disease | p.C431F | *Pathogenic* | 2 | *Pathogenic* |
| 10 | Disease and population | p.C441R | *Pathogenic* | 1 | *Pathogenic* |
| 11 | Disease | p.T240R | *Likely Pathogenic* | 2 | *Pathogenic* |
| 12 | Disease and population | p.R275W | *Likely pathogenic* | 1 | *Pathogenic* |
| 13 | Disease and population | p.G430D | *Likely pathogenic* | 2 | *Pathogenic* |
| 14 | Disease and population | p.A46T | *Likely benign* | 3 | *Likely benign* |
| 15 | Disease and population | p.P153R | *Likely benign* | 4 | *Benign* |
| 16 | Disease and population | p.M192L | *Likely benign* | 4 | *Benign* |
| 17 | Disease and population | p.R402C | *Likely benign* | 4 | *Benign* |
| 18 | Disease and population | p.Q34R | *Benign* | 3 | *Benign* |
| 19 | Disease and population | p.A82E | *Benign* | 4 | *Benign* |
| 20 | Disease and population | p.S167N | *Benign* | 4 | *Benign* |
| 21 | Disease and population | p.R334C | *Benign* | 5 | *Benign* |
| 22 | Disease and population | p.V380L | *Benign* | 4 | *Benign* |
| 23 | Disease and population | p.D394N | *Benign* | 4 | *Benign* |
| 24 | Disease and population | p.D18N | *Uncertain significance* | 4 | *Likely benign* |
| 25 | Disease and population | p.R33Q | *Uncertain significance* | 4 | *Likely benign* |
| 26 | Disease and population | p.P37L | *Uncertain significance* | 5 | *Likely benign* |
| 27 | Disease and population | p.R42C | *Uncertain significance* | 4 | *Likely benign* |
| 28 | Disease and population | p.R42H | *Uncertain significance* | 4 | *Likely benign* |
| 29 | Disease | p.W54R | *Uncertain significance* | 3 | *Uncertain significance* |
| 30 | Disease and population | p.R104W | *Uncertain significance* | 3 | *Uncertain significance* |
| 31 | Disease | p.K161N | *Uncertain significance* | 2 | *Pathogenic* |
| 32 | Disease | p.M192V | *Uncertain significance* | 4 | *Uncertain significance* |
| 33 | Population | p.V224A | *Uncertain significance* | 5 | *Likely benign* |
| 34 | Disease and population | p.R234Q | *Uncertain significance* | 5 | *Likely benign* |
| 35 | Disease and population | p.T240M | *Uncertain significance* | 2 | *Pathogenic* |
| 36 | Population | p.V248I | *Uncertain significance* | 4 | *Likely benign* |
| 37 | Disease | p.C253F | *Uncertain significance* | 1 | *Pathogenic* |
| 38 | Disease and population | p.R256C | *Uncertain significance* | 5 | *Likely benign* |
| 39 | Disease and population | p.D280N | *Uncertain significance* | 4 | *Likely benign* |
| 40 | Disease | p.C289G | *Uncertain significance* | 1 | *Pathogenic* |
| 41 | Disease and population | p.E310R | *Uncertain significance* | 4 | *Likely benign* |
| 42 | Disease | p.G328E | *Uncertain significance* | 4 | *Uncertain significance* |
| 43 | Disease and population | p.R334H | *Uncertain significance* | 4 | *Uncertain significance* |
| 44 | Disease | p.T351P | *Uncertain significance* | 4 | *Uncertain significance* |
| 45 | Disease and population | p.G359D | *Uncertain significance* | 3 | *Uncertain significance* |
| 46 | Disease and population | p.R366W | *Uncertain significance* | 4 | *Likely benign* |
| 47 | Disease and population | p.R396G | *Uncertain significance* | 4 | *Uncertain significance* |
| 48 | Disease | p.C418R | *Uncertain significance* | 1 | *Pathogenic* |
| 49 | Disease and population | p.P437L | *Uncertain significance* | 2 | *Pathogenic* |
| 50 | Disease and population | p.E444Q | *Uncertain significance* | 4 | *Uncertain significance* |
| 51 | Disease and population | p.M458L | *Uncertain significance* | 5 | *Likely benign* |

### *Parkin* variants can be classified functionally in cells

Next, to determine the effects of the variants on Parkin function, we employed a cell-based assay to monitor and quantify mitophagy using fluorescence-activated cell sorting (FACS) in U2OS cells stably expressing inducible mt-Keima, a pH-sensitive fluorescent protein that is targeted to mitochondria and exhibits a large shift in its emission wavelength upon engulfment in the acidic compartment of lysosomes (27). U2OS cells express very low level of endogenous Parkin, which is insufficient to mediate mitophagy in response to the mitochondrial potential uncoupler CCCP (28). In cells transiently expressing either wildtype (WT) Parkin or one of the Parkin variants fused to GFP, the shift in mt-Keima emission was measured upon four hours of mitochondrial depolarization with CCCP (Supplementary Fig. 3A). The GFP intensity in untreated cells was also quantified as a measure of steady-state Parkin protein levels (Supplementary Fig. 3B). The level of CCCP-induced Parkin-mediated mitophagy and GFP intensity of the Parkin variants were normalized to that of WT Parkin.

**Figure 3.**
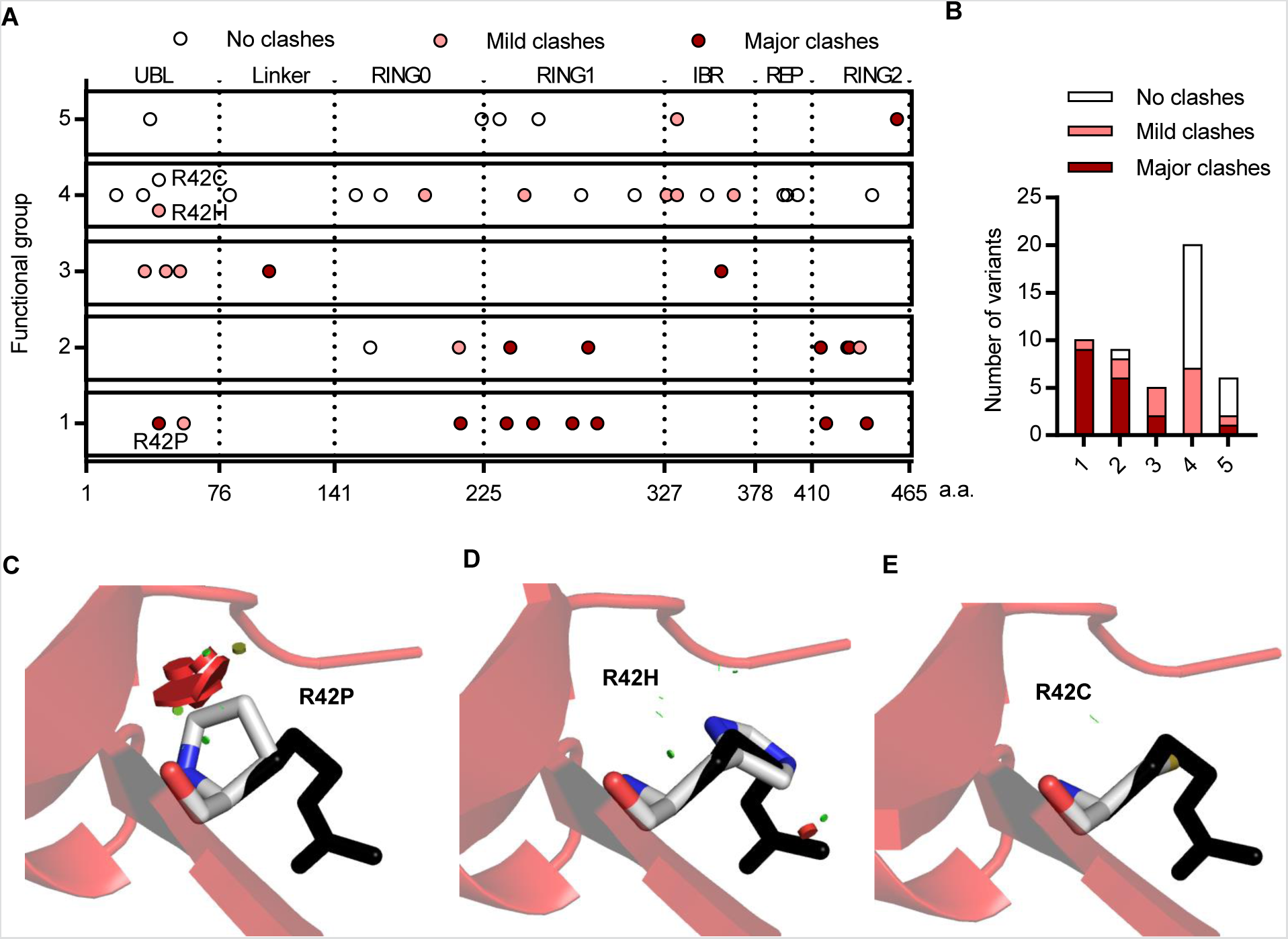
Steric clashes in structural simulations predicted the dysfunction of *Parkin* variants. **(A)** Schematic representation of Parkin missense variants on Parkin protein 2D structure. Each circle indicates a missense variant. The location of the variant on the 2D sequence was plotted in the X axis with dotted lines separating the Parkin domains. The functional groups described in Figure 2A, were plotted on the Y axis. The colors indicate the type of clash introduced by the missense variant from structural simulation. **(B)** Distribution of the variants from (A) within functional groups according to the type of clash they introduce. **(C)** Structure of human Parkin bound to phospho-ubiquitin (PDB 5n2w) was used to illustrate the impact of the R42P mutation. Substitution of the arginine side chain (black) to proline (white) introduced major clashes (red disks), which would destabilize the β-sheet in the Ubl. **(D)** Substitution of the arginine side chain (black) to histidine (white) introduced mild clashes (red and green disks). **(E)** Substitution of the arginine side chain (black) to cysteine (white) did not introduce any clashes.

We analyzed all the variants that were designated clinically as *pathogenic* or *likely pathogenic*, *benign* or *likely benign* (Fig. 1B). We also analyzed an additional twenty-eight variants of *uncertain significance*, including variants that were reported as homozygous or compound heterozygous in either the disease-specific databases or in population databases (Fig. 1C and Supplementary Table 1). These variants represented the most common missense variants reported in the human population from public databases. Moreover, they were reported in 283 out of 309 families or isolated patients carrying Parkin missense variants in PD-specific databases.

The thirteen *Parkin* variants classified as *pathogenic* or *likely pathogenic* based on clinical evidence all showed significantly decreased mitophagy (Fig. 1B). Seven variants (p.R42P, p.V56E, p.C212Y, p.C253Y, p.C238W, p.R275W, and p.C441R) also showed decreased GFP intensity (Fig. 1B), suggesting reduced protein stability. Of these, all the variants in the R0RBR of Parkin formed inclusions to different degrees, detected by fluorescence microscopy (Supplementary Fig. 4). In contrast, the p.R42P and p.V56E variants in the Ubl domain showed lower overall GFP intensity without visible inclusions. Thus, for a subset of *pathogenic* and *likely pathogenic Parkin* variants, the observed defects in mitophagy are likely to stem from abnormal protein folding and reduced protein stability. In contrast, all ten *Parkin* variants classified as *benign* and *likely benign* based on clinical evidence exhibited similar GFP intensity as WT and most, with three exceptions, also exhibited WT levels of Parkin-mediated mitophagy (Fig. 1B). p.Q34R and p.A46T displayed decreased mitophagy, whereas p.R334C showed an almost three-fold increase (Fig. 1B). Of the twenty-eight variants of *uncertain significance* based on clinical evidence, fourteen displayed similar GFP intensity and mitophagy as WT Parkin (Fig. 1C). Nine displayed impaired mitophagy, three of which also showed decreased GFP intensity. Surprisingly, five variants showed increased mitophagy (Fig. 1C).

**Figure 4.**
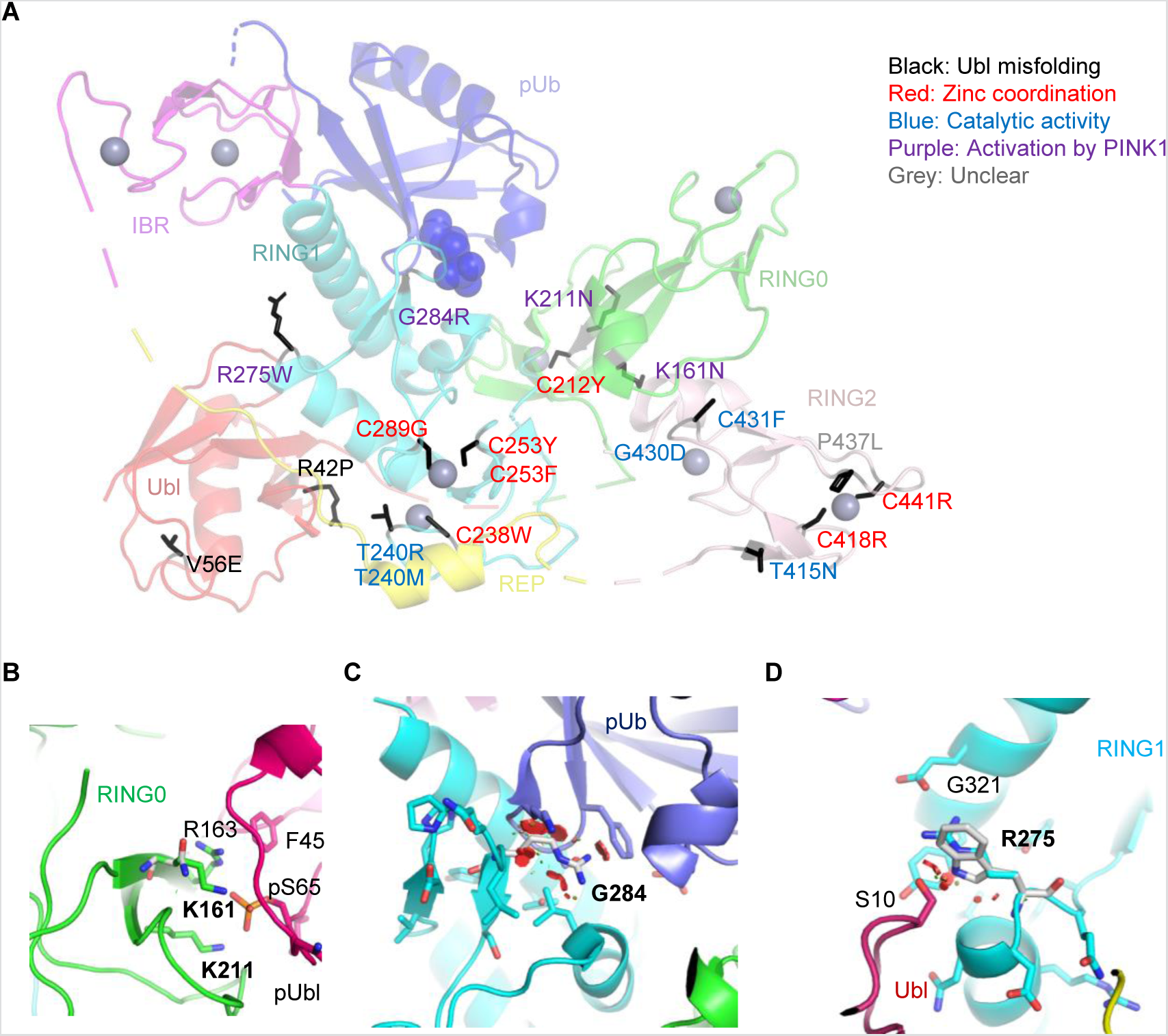
Structural analysis of *Parkin* variants revealed various pathogenic mechanisms. **(A)** Pathogenic variants were mapped onto the 3D structure of human Parkin bound to phospho-ubiquitin (PDB 5N2W). The side-chains of the amino acids substituted by the variants were highlighted in black. The blue spheres represent the phosphate of pUb. The grey spheres represent zinc. The color of the text indicates the type of disruption to Parkin caused by the variant. **(B)** Close-up view of pUbl-RING0 interface in the structure of fly pParkin bound to phospho-Ub (PDB 6DJX). Lys161 and Lys211 form ionic interactions with the phosphate on Ser65 of the pUbl. Mutations of these lysine residues would weaken the pUbl-RING0 interaction, preventing activation of Parkin. **(C)** Close-up view of the pUb:RING1 interface in human Parkin bound to pUb (PDB 5N2W). The G284R variant in Parkin RING1 would introduce major clashes with pUb, disrupting the interaction. **(D)** Close-up view of Arg275 in human Parkin bound to pUb (PDB 5N2W). Arg275 interacts with Glu321 in the helix that interacts with pUb. Mutation to a tryptophan (white) would introduce clashes with this helix as well as Ser10 in the Ubl domain.

The wide range of changes in Parkin levels and mitophagy prompted us to ask whether we could classify the different variants into discrete groups. Variants that significantly decreased Parkin protein levels and mitophagy were assigned to Group 1 (Fig. 2A and Table 1). Variants that severely (0-30% of WT) or moderately (30-60% of WT) reduced Parkin-mediated mitophagy but displayed normal protein levels were assigned to Groups 2 and 3, respectively. Group 4 consisted of variants that were similar to WT, whereas Group 5 consisted of variants with increased (>140% of WT) Parkin-mediated mitophagy. Interestingly, all the *pathogenic* or *likely pathogenic* variants, classified based on clinical evidence, were assigned to group 1 or 2, whereas all the *benign* or *likely benign* variants fell into group 3, 4, or 5 (Fig. 2A-B). This suggested that the two functional measurements in our cell model could faithfully discriminate the pathogenic variants from the “non-pathogenic” variants.

### Integration of clinical and functional evidence refines the classification of *Parkin* variants

We next examined which clinical features were most strongly correlated with functional alterations. Among the variants that segregated with PD in families (Supplementary Table 1), all but one (p.R33Q) severely altered Parkin function and were assigned to groups 1 or 2 (Fig. 2C). However, it is important to note that segregation analysis was only possible in ~30 families, as most reports only involved single case reports without related family information (Supplementary Table 1). Conversely, variants that were reported as homozygotes in ExAC were frequent in Groups 3, 4 and 5 and not assigned to Groups 1 or 2 (Fig. 2D). Notably, several variants that were reported in PD patients as homozygous or compound heterozygous, nonetheless displayed WT Parkin levels and function (Fig. 2E). Most had relatively high MAFs in ExAC (Supplementary Table 1), suggesting their presence in patients was due to their high prevalence rather than pathogenicity. Taken together, our data shows that variants segregating with disease in families impair Parkin function (Groups 1 and 2), whereas variants that occur as homozygotes in ExAC did not severely reduce Parkin function (Groups 3, 4 and 5).

Next, we devised a scoring scheme, based on Sherloc, to assign benign or pathogenic points (Supplementary Fig. 5), according to the functional group to which the *Parkin* variants were assigned in the cellular assays (Fig. 2A). The pathogenic points and benign points were then added to the corresponding points from the clinical evidence in order to obtain combined *pathogenic* and *benign* scores from all evidence for the final annotation of each variant (Table 1 and Supplementary Table 1). Using these combined clinical and functional scores, all the *likely pathogenic* variants from clinical evidence were reclassified as *pathogenic* (Table 1). Additionally, three *likely benign* variants were reclassified as *benign* (Table 1). Most of the variants of *uncertain significance* were reclassified as *likely benign*, while six were reclassified as *pathogenic* and nine remained of *uncertain significance* (Table 1). In summary, all variants that caused functional alterations assigned to groups 1 and 2, as measured in our experimental assay, were reclassified as *pathogenic,* whereas most variants assigned to functional groups 3, 4 and 5 were reclassified as either *benign* or *likely benign* (Fig. 2F).

**Figure 5.**
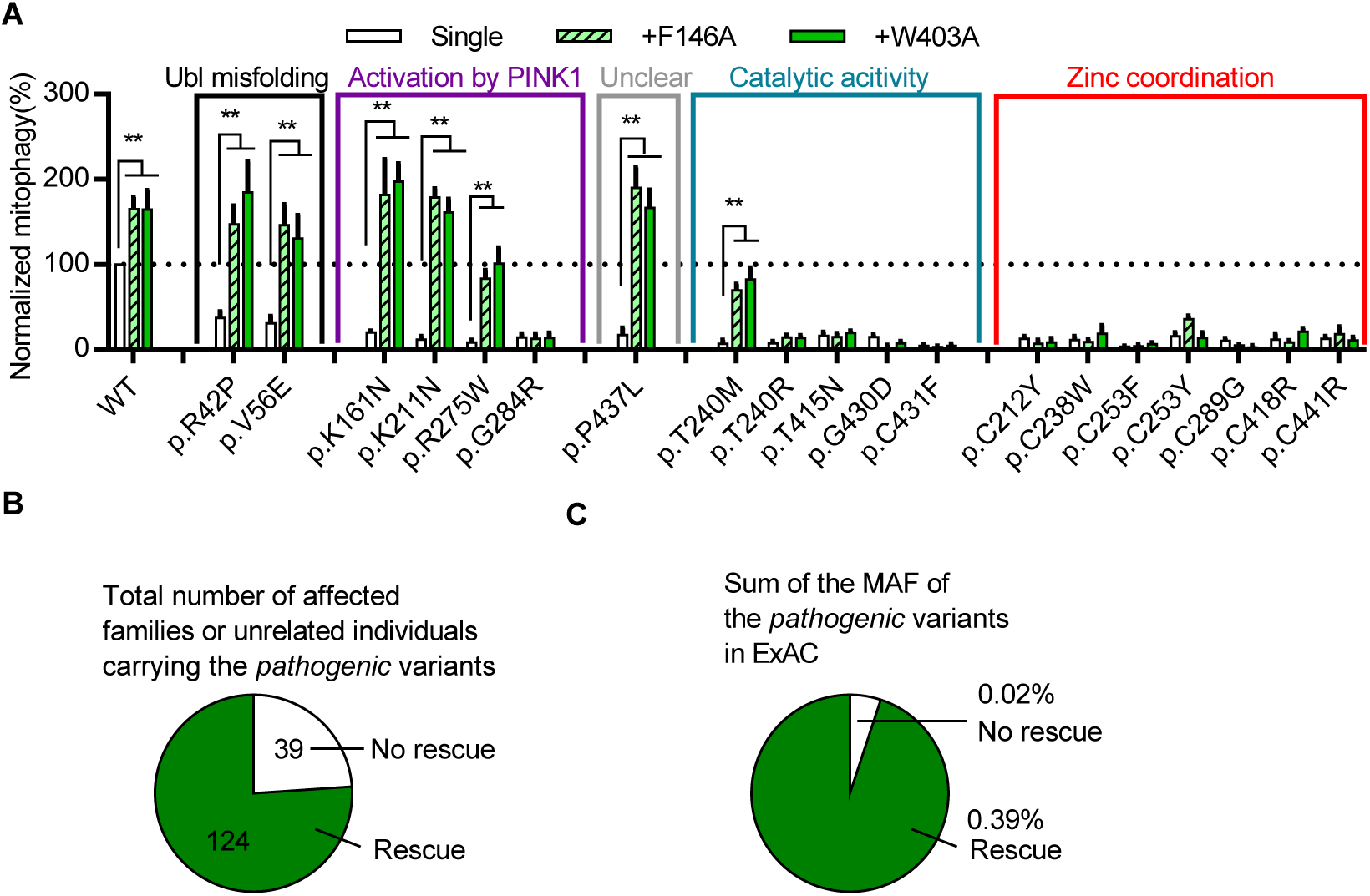
Structure-guided designer hyperactive Parkin mutants can rescue mitophagy in *pathogenic* variants. **(A)** Quantification of induced mitophagy after 4h of CCCP treatment in U2OS cells expressing WT GFP-Parkin, *pathogenic* missense variants, or W403A or F146A *in cis* with WT or *pathogenic* variants. Mitophagy mediated by each Parkin missense variant was normalized to that of WT Parkin in each replicate. * P<0.05, ** P<0.01, in two-way ANOVA with Dunnett’s post-hoc test comparing the function of each variant with the variant *in cis* with W403A or F146A. N=3-7. **(B)** The number of families or individuals with PD carrying the *pathogenic* missense variants for which mitophagy was or was not rescued by the designer mutations are shown. **(C)** The sum of the MAF in ExAC of the *pathogenic* missense variants for which mitophagy was or was not rescued by the designer mutations are shown.

### Structural analysis of *Parkin* variants reveals pathogenic mechanisms

As the structure of Parkin is known, we modeled the effects of variants on the reported crystal structures of Parkin to gain insight into the mechanisms underlying the functional changes. The structures of autoinhibited Parkin, pUb-bound Parkin, p-Ub-bound phospho-Parkin, pUb-E2 enzyme-bound phospho-Parkin were used as they depict Parkin in its different states of activation (17, 18, 29, 30).

Most variants in functional groups 1 and 2 were predicted to introduce steric clashes with nearby residues in at least one of the Parkin structures. In contrast, most variants in Group 4, which displayed similar function as WT Parkin, introduced no major clashes and did not affect interactions (Fig. 3A-B). However, mild clashes were observed in a few cases, suggesting that some degree of steric clashing could be tolerated and possibly be compensated for by local conformation changes to maintain overall protein function. For example, three variants at Arg42 were analyzed. Mutation of Arg42 to proline introduced major clashes, whereas substitution to histidine introduced mild clashes and there was no clash introduced by substitution to cysteine (Fig. 3C-E). Congruently, only p.R42P was classified as *pathogenic* based on clinical criteria and functional impairment in cell-based assays (Fig. 1B-C), consistent with the simulations that showed this mutation unfolded the Ubl domain (22, 31). These simulated steric clashes nicely illustrate how different amino acid substitutions at a given residue could lead to distinct functional impairment.

The effects of variants in Groups 1 and 2 likely disrupt several different aspects of Parkin function (Fig. 4A). Seven variants involve cysteine residues that coordinate zinc, and their mutation would result in overall misfolding of the Parkin protein (29). Five variants alter key motifs mediating ubiquitination of substrates, including steric clashes with the E2 binding site on RING1 (p.T240R and p.T240M), and residues implicated in thioester transfer of Ub in the catalytic RING2 domain (p.T415N, p.G430D, p.C431F) (32). These types of alterations are likely to cause a complete loss of Parkin function in either PINK1-Parkin mediated mitophagy or other potential Parkin pathways. Four variants specifically localize to motifs implicated in the conformational change that occurs during activation by PINK1. p.K161N and p.K211N introduce no or mild steric clashes, but substitution of the basic lysine residue to the neutral asparagine eliminates the interaction with the acidic phosphate in pUbl (Fig. 4B and (17, 18)). p.G284R introduces major clashes with pUb, thus impairing binding and recruitment to mitochondria (Fig. 4C). p.R275W disrupts interaction with the helix that mediates pUb binding (Fig. 4D). This is predicted to destabilize Parkin, consistent with the observed decreased steady-state protein level and the presence of cellular inclusions (Fig. 1B and Supplementary Fig. 4). This helix becomes more exposed following pUb binding and may also be involved in the allosteric release of the Ubl during activation (Fig. 4D and (17, 18)). The clashes caused by p.R42P and p.V56E are predicted to misfold the Ubl domain and thus destabilize the protein, as demonstrated earlier for p.R42P (22). Concurrently, p.R42P and p.V56E may also hinder conformational change during activation by preventing Ubl phosphorylation (33) and the binding of pUbl to RING0 (17). One variant, p.P437L, introduced very mild steric clashes in RING2 in the Parkin structure, and the exact molecular mechanism of p.P437L in causing decreased mitophagy remains unclear.

Several variants in functional Group 3 that moderately decreased Parkin-mediated mitophagy might also act by hindering Parkin activation by PINK1. p.A46T could disrupt the interaction of pUbl with RING0 (Supplementary Fig. 6A). p.R104W induced clashes to the newly identified activation element (ACT), which binds RING0 and helps stabilize the interaction with pUbl and RING0 (Supplementary Fig. 6B and (18)). p.G359D disrupted the glycine-rich loop in the IBR domain that interacts with pUb (Supplementary Fig. 6C). These clashes may be compensated by minor local conformation changes, and thus lead to milder disruptions in mitophagy and protein stability.

**Figure 6.**
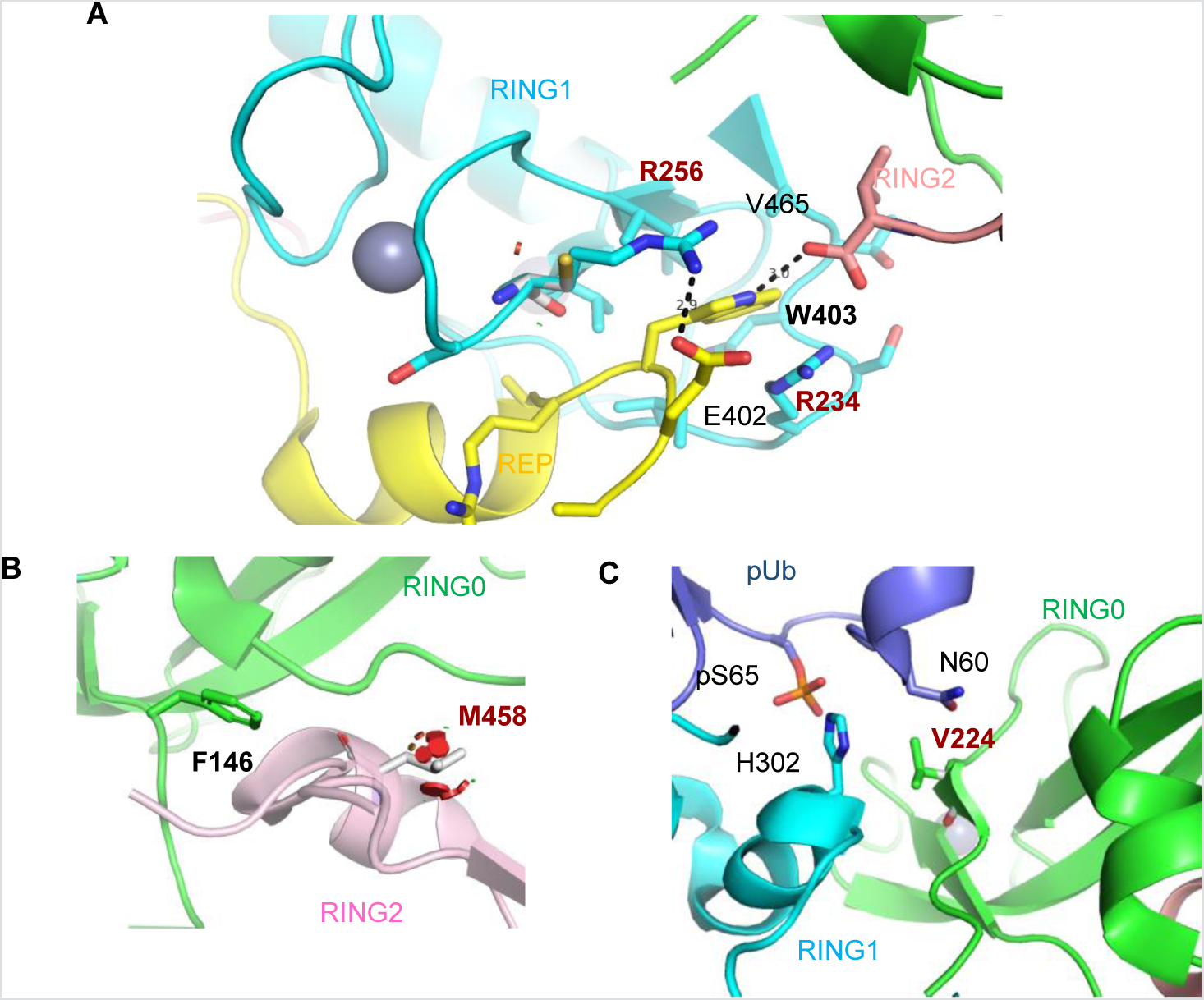
Structural basis for the effects of the naturally occurring hyperactive Parkin variants. **(A)** Close-up view of R234Q and R256C variant sites in the structure of human Parkin bound to pUb (PDB 5N2W). Arg256 forms a hydrogen bond with Glu402, and its mutation to a cysteine would destabilize the REP:RING1 interaction, similar to W403A. The side-chain of Arg234 also stacks with the indole ring of Trp403. **(B)** Close-up view of M458L variant site in the RING0:RING2 interface (PDB 5N2W). M458L introduced major clashes that would destabilize the RING0:RING2 interaction, similarly to F146A. **(C)** Close-up view of V224A variant site (PDB 5N2W). Val224 interacts with pUb and forms van der Waals force interactions with Asn60. Mutation to alanine could modulate the affinity for pUb.

### Structure-guided *designer* hyperactive Parkin mutants can rescue mitophagy in pathogenic variants

We next hypothesized that some of the *pathogenic* variants may represent targets for genotype-specific therapy. As a proof of concept, we introduced two artificially designed mutations, W403A and F146A, which destabilized the REP (repressor element of Parkin):RING1 and the RING0:RING2 interfaces, respectively. Both mutations have been shown to accelerate mitophagy by promoting the conformational changes that occur at these interfaces during Parkin activation by PINK1 (28, 29). We tested whether these *hyperactive* mutations could rescue the mitophagy defects seen in the *pathogenic* Parkin variants. As reported previously, mutating F146A or W403A alone enhanced mitophagy compared to WT Parkin (Fig. 5A) (28). Remarkably, introducing F146A or W403A *in cis* with p.R42P, p.V56E, p.K161N, p.K211N, p.R275W p.P437L or p.T240M rescued mitophagy (Fig. 5A). Variants p.R42P, p.V56E and p.R275W each lower Parkin levels, likely by disrupting protein stability (Fig. 1B). However, introduction of F146A or W403A did not restore Parkin levels to WT (Supplementary Fig. 7), suggesting the rescue of mitophagy was mediated by Parkin activation *per se* rather than by enhancing Parkin protein stability. The *pathogenic* variants p.K161N and p.K211N are involved in binding the pUbl during activation (17, 18). The fact that both these variants can be rescued by F146A or W403A suggests that destabilization of the REP:RING1 or the RING0:RING2 interface can bypass the tethering of the Ubl to RING0, which occurs during Parkin activation.

**Figure 7.**
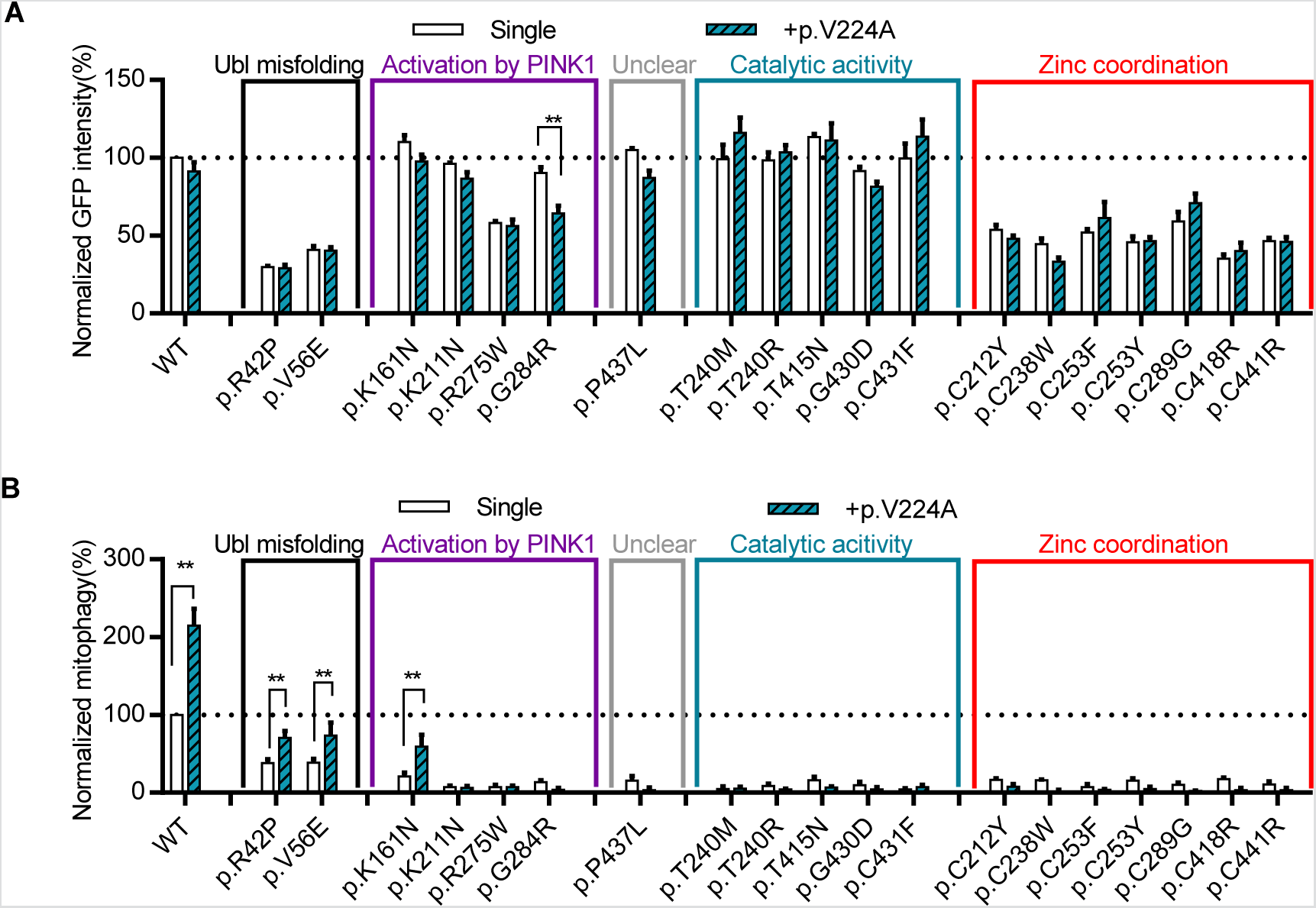
The naturally-occurring *Parkin* p.V224A hyperactive variant rescued mitophagy in several *pathogenic* variants. **(A)** Quantification of GFP intensity from GFP signal by FACS in untreated cells expressing WT GFP-Parkin, *pathogenic* missense variants, or p.V224A *in cis* with WT or *pathogenic* variants. The GFP intensity for each Parkin missense variant was normalized to that for WT Parkin in each replicate. **(B)** Quantification of induced mitophagy after 4h of CCCP treatment in cells expressing WT GFP-Parkin, *pathogenic* missense variants, or p.V224A *in cis* with WT or *pathogenic* variants. Mitophagy mediated by each Parkin missense variant was normalized to that of WT Parkin in each replicate. * P<0.05; ** P<0.01, in two-way ANOVA with Dunnett’s post-hoc test comparing the function of each variant with variant *in cis* with p.V224A. N=3-7.

Introducing F146A or W403A *in cis* with the variants could not rescue any of the seven *pathogenic* cysteines variants involved in zinc coordination (Fig. 5A). Thus, the severe disruption of Parkin folding and stability induced by mutating these cysteines are likely to preclude them from being good candidates for therapeutic rescue by hyperactivation (Fig. 1B-C). Similarly, most variants that disrupted key catalytic sites could not be rescued (Fig. 5A). Also, p.G284R could not be rescued as it disrupts binding to pUb, which is an essential receptor for recruiting Parkin to damaged mitochondria (34). Interestingly, whereas both p.T240M and p.T240R are predicted to interfere with E2 Ub-conjugating enzyme binding to RING1 and showed similar severe defects in mitophagy, only the former could be rescued by F146A or W403A (Fig. 5A). Both variants created clashes in the E2 binding site (Supplementary Fig. 8A-B). However, Arg240 created a positive charge at the interface, increasing the electrostatic repulsion of E2. Methionine is less bulky and neutral, and its flexibility could allow some weak interactions with E2 to remain, perhaps explaining its rescue by F146A or W403A (Supplementary Fig. 8C-E).

Overall, the defects in mitophagy of seven of the 19 *pathogenic* variants could be rescued by the designed activating mutations. These seven variants are responsible for over 75% of reported PD patients carrying *pathogenic* missense variants (Fig. 5B) and were the most frequent *pathogenic* missense variants in the general population (Fig. 5C). Mimicking the effects of F146A or W403A could therefore be a useful starting point for designing treatments for patients with PD caused by these *Parkin* variants.

### Characterization of naturally-occurring hyperactive Parkin variants

We identified six naturally-occurring variants that, considering all the evidence, were classified as *likely benign* or *benign* and showed enhanced Parkin-mediated mitophagy (Fig. 1B, 1C and 2G). The Parkin structure shows that Arg234 and Arg256 are located at the interface between the REP and RING0. The p.R234Q and p.R256C variants are predicted to destabilize the interface, thus mimicking the W403A *designer* mutant used above (Fig. 6A). Additionally, p.M458L may destabilize the RING0:RING2 interface, mimicking the effects of our other *designer* mutation, F146A (Fig. 6B). Thus, based on structural predictions, three of the six naturally-occurring variants are likely to activate Parkin via mechanisms akin to those involved in the hyperactive mutants designed previously (28). Because these variants occur naturally in the population, our findings demonstrating rescue of mitophagy suggest that targeting these sites and mechanisms are likely to be tolerated and potentially therapeutic in PD. p.P37L also moderately increased Parkin-mediated mitophagy, but the structural basis of the increased activity was unclear, as this variant does not create any steric clash and thus should not affect interactions of the Ubl with RING1 or interactions of the pUbl with RING0 (Supplementary Fig. 9A-B). Mutation of Arg334 to a cysteine could affect the coordination of a nearby zinc in the IBR, which may stabilize the interaction with pUb and thereby enhance Parkin activity (Supplementary Fig. 9C).

Unlike the five other naturally-occurring hyperactive variants, p.V224A has not been reported in PD patients (Supplementary Table 1) and showed the highest (almost 3-fold above WT) Parkin-mediated mitophagy activity (Fig. 1C). The Val224 residue is localized in the pUb binding pocket, with its side-chain facing towards the phosphorylated Ser65 residue of pUb, and the mutation to alanine could modulate the affinity of Parkin for phospho-ubiquitin (Fig. 6C). We therefore examined the ability of hyperactive p.V224A to rescue the function of the *pathogenic* variants. Introducing the p.V224A variant *in cis* did not affect the protein level of most *pathogenic* variants, except for p.G284R (Fig. 7A). This may stem from an additive destabilizing effect of the double mutant on Parkin folding as both p.V224A and p.G284R are located within the same pUb binding motif. Introducing the p.V224A variant partially rescued the mitophagy defects of p.R42P, p.V56E and p.K161N (Fig. 7B). How the predicted effects of V224A on pUb-binding could partially compensate for defects in Ubl- and pUbl-mediated activation by p.R42P, p.V56E and p.K161N remains to be elucidated. p.V224A could not rescue the mitophagy deficit in p.R275W and p.G284R variants, nor could it rescue any of the remaining *pathogenic* variants that directly damaged catalytic activity and zinc coordination (Fig. 7B). Compared with F146A or W403A, p.V224A was less effective at rescuing the *pathogenic* variants, suggesting that the pUb-binding site may be a less promising target for activating mitophagy than releasing the autoinhibited conformation of Parkin.

## Discussion

*Parkin* mutations are the most common cause of recessive early-onset PD (EOPD) (2). Although *Parkin* loss-of-function is well established in EOPD (5, 35), causality for any given missense variant has been more difficult to ascertain. In this study, we have integrated clinical, experimental and structural modeling approaches to map out the landscape of *Parkin* variants in the general population and in PD patients. Our hope is that this work will help provide a more cohesive framework to guide basic science studies exploring the molecular and cellular functions of the *PINK1/Parkin* pathway and to guide clinicians caring for patients carrying specific *Parkin* variants. We also hope that the work will inform structure-based drug design to develop Parkin activators and help guide Parkin allele- and genotype-specific clinical studies.

For the over 200 *Parkin* variants reported in public databases (3, 5, 8), we found that only a minority of variants could be clearly designated as *likely pathogenic*, *pathogenic*, *likely benign* or *benign* based on clinical evidence alone. While this may not seem surprising for variants found only in population databases such as ExAC, where accompanying clinical information is scant, we found a similar situation for variants reported in patients. Indeed, 52 out 75 variants reported in PDmutDB and MDSgene remained of *uncertain significance* after having been subjected to the Sherloc algorithm, the variant classification framework derived from ACMG standards that we used in this study. The overarching message from these observations is that the mere presence of *Parkin* variants in PD patients, PD kindreds or PD-specific databases should be interpreted with caution and not automatically taken to imply pathogenicity. Rather, we propose that clinical evidence available for new variants should be subjected to the same rigorous classification scheme presented here to determine pathogenicity.

In addition to analyzing clinical evidence, we extensively characterized the cellular effects of the 51 *Parkin* variants most commonly found in patient and population sequencing databases. To our knowledge, a systematic analysis integrating clinical evidence with cellular function, on this scale, has not been reported previously for *Parkin* (20-25, 36, 37). Notably, all the variants designated as *pathogenic* or *likely pathogenic* based on clinical evidence also displayed severe mitophagy defects in cells (functional groups 1 and 2). Conversely, all variants designated clinically as *benign* or *likely benign* displayed mitophagy function in the WT range or showed only a slight reduction (functional groups 3, 4 and 5). While this may seem *a priori* as self-evident, several alternative functions of Parkin in cells have been proposed and the role of mitophagy in PD has yet to be definitively established (38-41). Thus, while this work does not refute the biological importance of such alternative functions, the tight correlation between the clinical impact of the variants and their effects on mitophagy provides further evidence that mitophagy can be used as a robust and disease-relevant readout of Parkin function that likely reflects a key pathogenic process in PD.

Assignment of the *Parkin* variants to functional groups in cells allowed us to determine which clinical features best correlate with and could be used to predict pathogenicity. Segregation of variants with PD in families and observation of homozygotes in ExAC turned out to be very strong predictors for pathogenicity or the absence of pathogenicity, respectively. In contrast, the mere report of PD patients with one or two *Parkin* variants or the absence of these variants in control cohorts or population databases should not be automatically taken to imply pathogenicity. Perhaps more importantly, integrating clinical with functional evidence from cells allowed us to re-assign 19 of the 28 variants, designated as of *uncertain significance* based on clinical evidence alone, to one of the benign and pathogenic categories. It also allowed us to reclassify 6 of the “*likely*” variants to their respective more definitive *benign* and *pathogenic* categories. Together, these findings attest to the power of using this sort of iterative combined clinical and experimental approach to stratify variants.

Given what is already known about the structure and function of the Parkin protein, the work enables in-depth mechanistic exploration of how *pathogenic* variants can lead to dysfunction. This is something that has been sorely lacking and may have important implications for how best to target Parkin and design activators for future therapy. For most of the *pathogenic* variants, the mechanisms by which they interfere with function can be rationalized based on the Parkin structure. As an important proof of concept, we showed that the function of several pathogenic *Parkin* variants, defective in mitophagy in cells, could be rescued when expressed *in cis* with mutations that have been previously designed to enhance Parkin activity (28, 29). This provides further stratification according to therapeutic potential. For instance, alterations in residues involved in zinc coordination, in catalytic activation or in pUb-binding could not be rescued. In contrast, alterations in residues involved in Ubl folding or in the pUbl-RING0 interface in the active Parkin structure could be fully rescued, suggesting that therapeutics that disrupt the REP-RING1 or the RING0-RING2 interfaces could potentially bypass these defects (17, 18). Importantly, many of the most commonly occurring variants were the ones that could be rescued, something that bodes well for patients carrying these variants, should a therapeutic mimicking W403A or F146A become available.

One of the most surprising findings of our study was that several naturally-occurring variants exhibited a 1.5- to almost 3-fold enhancement in Parkin-mediated mitophagy in our assay. This could not simply be explained by increased Parkin protein levels or by the fact that our assay involved overexpression. Indeed, except for certain pathogenic variants that destabilized Parkin, most variants, including the hyperactive ones, displayed steady-state levels that were very close to WT levels. These hyperactive variants provide an important proof of principle that there are, presumably healthy, individuals in the population living with enhanced Parkin activity. The strongest activating variant was V224A, which increases mitophagy by nearly 3-fold and occurs very near the site for pUb-binding. This was surprising as pUb-binding serves as a critical receptor to recruit Parkin to mitochondria and, to date, every reported mutation in this motif, abolished or dramatically reduced mitophagy. Moreover, when expressed *in cis*, V224A partially rescued certain, but not all, of the mutants that were rescued by W403A and F146A. In the future, it will be important to test whether this involves an enhancement in pUb-binding or some other downstream allosteric effect. Similarly, it will be important to determine the mechanisms by which the two remaining hyperactive variants, P37L and R334C, enhance Parkin function. Moreover, as we only sampled 51 of the over 200 *Parkin* variants in the population in this study, it is conceivable that other yet-to-be-discovered hyperactive variants will provide further mechanistic insights into Parkin activation and help identify additional therapeutic sites within the protein.

## Materials and methods

### Classification of *Parkin* missense variants

We utilized Sherloc (semiquantitative, hierarchical evidence-based rules for locus interpretation), a variant classification framework derived from ACMG standards to assign *Parkin* missense variants into five categories: *pathogenic*, *likely pathogenic*, *benign*, *likely benign* and of *uncertain significance* (9). We considered two broad categories of evidence for the classification, clinical and functional. The procedures for evaluating and scoring these lines of evidence are summarized as root-decision trees in Supplementary Fig. 1, 2, and 5.

For clinical evidence, information regarding missense variants in *Parkin* reported in the population database ExAC (http://exac.broadinstitute.org/gene/ENSG00000185345) (8) and the disease-specific databases, PDmutDB (http://www.molgen.vib-ua.be/PDmutDB) (3), and MDSgene (http://www.mdsgene.org/) (5) were searched. We also searched dbSNP (http://www.ncbi.nlm.nih.gov/snp) and the Exon variant server (EVS, http://evs.gs.washington.edu/EVS/) for missense variants that were found in the disease databases, but not in ExAC. The homozygote count, MAF in ExAC and maximal MAF in dbSNP, EVS and subpopulations in ExAC were calculated and used to assign points to the variants according to the decision tree in Supplementary Fig. 1A. The clinical cases reported in PDmutDB and MDSmutDB were evaluated according to the decision tree in Supplementary Fig. 1B-C. The original references were traced back for the indexed families or individuals reported in these databases. Indexed cases reported in both databases cited from the same reference were only evaluated once. Indexed cases reported in a more recent paper showing the same information (same number of family members with same genotype and phenotype) as a case in an older reference were considered as the same family and the older report was used.

For functional evidence, the effects of the variants in the cellular assay were assigned points according to the decision tree in Supplementary Fig. 5. We imposed a 2.5-point cap on functional evidence to ensure that functional data which lacked supporting clinical evidence would not be sufficient on its own to reach the threshold required (3 benign points or 4 pathogenic points; Supplementary Fig. 2) to assign a variant to the *pathogenic* or *benign* categories (9).

### Cell culture, cloning and mutagenesis

Human osteosarcoma U2OS cells were a gift from Dr. Robert Screaton (Sunnybrook Research Institute). U2OS cells stably expressing mtKeima (a gift from A. Miyawaki, Laboratory for Cell Function and Dynamics, Brain Science Institute, RIKEN, Japan) were created by transfecting plasmid DNA using jetPRIME (Polyplus), followed by selection with G418 for 2 weeks and sorting using flow cytometry (28). Cells were maintained in DMEM supplemented with 10% fetal bovine serum (FBS), 4 mM L-glutamine and 0.1% Penicillin/Streptomycin, in a 37℃ incubator with 5% CO_2_. All GFP-Parkin variants were generated using PCR mutagenesis on the GFP-Parkin WT plasmid (addgene#45875) according to the manufacturer’s protocol (Agilent Technologies). Constructs were verified by Sanger sequencing.

### Mitophagy and GFP-intensity measurement by FACS

U2OS cells stably expressing ecdysone-inducible mt-Keima were induced with 10 mM ponasterone A and transiently transfected with WT or variant GFP-Parkin for 24 h and treated with DMSO or 20 μM CCCP for 4 h and followed immediately by flow cytometry. To minimize transfection efficiency variation, the same amount of GFP-Parkin WT or variant plasmid was utilized and only the population of GFP-positive cells were analyzed in the subsequent FACS data processing. For flow cytometry, cells were trypsin digested, washed and resuspended in PBS prior to their analysis on an LSR Fortessa (BD Bioscience) equipped with 405 and 561 nm lasers and 610/20 filters (Department of Microbiology and Immunology Flow Cytometry Facility, McGill University). Measurement of mtKeima was made using a dual-excitation ratiometric pH calculation where pH 7 was detected through the excitation at 405 nm and pH 4 at 561 nm (28). For each untreated sample, 75,000 events were collected and single GFP-Parkin-positive cells were subsequently gated for quantification of the geometric mean of the GFP signal as a measure of steady-state Parkin protein levels. The value for each Parkin missense variant was normalized to that for the WT in each experiment. For each untreated and treated sample, single GFP-Parkin-positive, mtKeima-405 nm-positive cells were subsequently gated. The percentage of cells with an increase in the 405nm:561nm ratio in mtKeima was quantified. The percentage in treated cells minus the percentage in untreated cells was calculated as the induced Parkin-mediated mitophagy. The induced mitophagy for each Parkin missense variant was normalized to that for WT in each repeat. Data was analyzed using FlowJo v10.1 (Tree Star).

### Modeling of Parkin Structures, Modifications and Variants

The structures of human Parkin bound to phospho-ubiquitin (PDB 5N2W), rat parkin (PDB 4ZYN), human phospho-parkin bound to phospho-Ub (PDB 6GLC) and fly pParkin-pUb-UbcH7 complex (PDB 6DJX) were analyzed using PyMOL version 1.5 (Schrödinger, New York). Mutations and clashes were simulated using the mutagenesis wizard toolbox. The presence of more than three simulated significant clashes (red disks) was taken to indicate major clashes. One to three significant clashes (red disks) together with other slight clashes (brown and green disks) were considered as minor clashes. Polar contacts within 4 Å distance of the residue were explored for characterizing interactions.

### Statistical analysis

For statistical analysis of mitophagy and protein levels, one-way analysis of variance (ANOVA) with Bonferroni post-hoc tests were performed. To determine the ability of hyperactive mutants to rescue *Parkin* variants, two-way analysis of variance (ANOVA) and Bonferroni post-hoc test comparing row factors among the single or double mutations were performed. *P<0.05; **P<0.01; ***P<0.001.

## Author contributions

W.Y. performed cloning, genetic analysis, and experiments in cells. E.J.M., M.Y.T. and A.I.K. assisted with cloning and experiments in cells. Z.G-O. assisted with genetic analysis. J.-F.T. assisted with all the structural simulations. W.Y., E.J.M., M.Y.T., Z.G-O, J.F.T. and E.A.F. participated in the design of experiments, data analysis and preparation of the manuscript.

## Acknowledgements

We thank members from the Trempe and Fon labs, as well as Kalle Gehring for useful discussion and comments. The flow cytometry work/cell sorting was performed in the McGill Life Science Complex Flow Cytometry Core Facility supported by funding from the Canadian Foundation for Innovation. We acknowledge support from Parkinson Society Canada (Basic Science Postdoctoral Fellowship to W.Y.), and the Canadian Institutes of Health Research (FDN grant – 154301 to E.A.F.).

