## Supplemental figures for "The Landscape of *Parkin* Variants Reveals Pathogenic Mechanisms and Therapeutic Targets in Parkinson’s Disease"

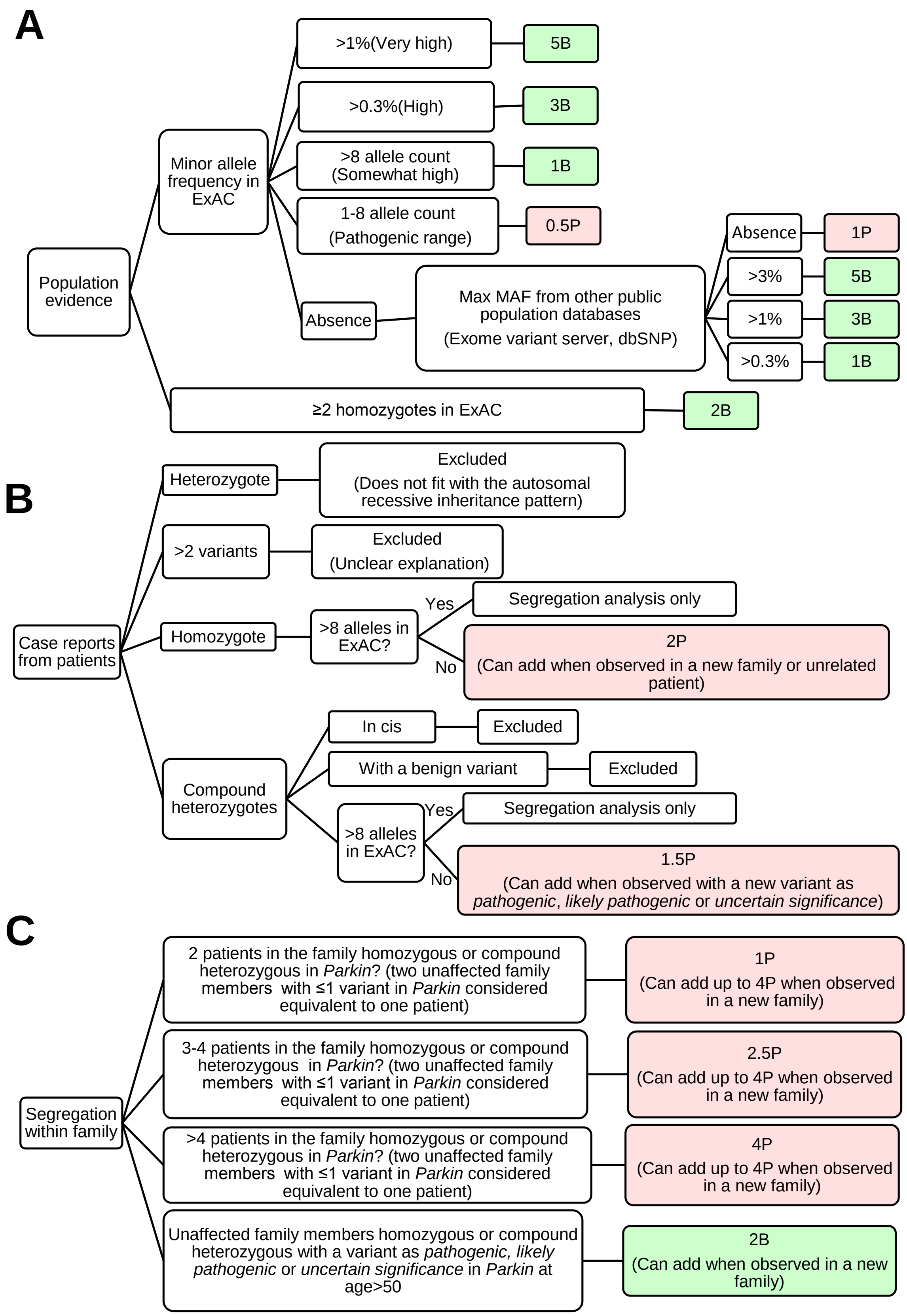
A
Population evidence
Minor allele frequency in ExAC
>1%(Very high)
5B
>0.3%(High)
3B
>8 allele count (Somewhat high)
1B
1-8 allele count (Pathogenic range)
0.5P
Absence
Max MAF from other public population databases (Exome variant server, dbSNP)
Absence
1P
>3%
5B
>1%
3B
>0.3%
1B
≥2 homozygotes in ExAC
2B
B
Case reports from patients
Heterozygote
Excluded (Does not fit with the autosomal recessive inheritance pattern)
>2 variants
Excluded (Unclear explanation)
Homozygote
>8 alleles in ExAC?
Yes
Segregation analysis only
No
2P (Can add when observed in a new family or unrelated patient)
Compound heterozygotes
In cis
Excluded
With a benign variant
Excluded
>8 alleles in ExAC?
Yes
Segregation analysis only
No
1.5P (Can add when observed with a new variant as *pathogenic*, *likely pathogenic* or *uncertain significance*)
C
Segregation within family
2 patients in the family homozygous or compound heterozygous in *Parkin*? (two unaffected family members with ≤1 variant in *Parkin* considered equivalent to one patient)
1P (Can add up to 4P when observed in a new family)
3-4 patients in the family homozygous or compound heterozygous in *Parkin*? (two unaffected family members with ≤1 variant in *Parkin* considered equivalent to one patient)
2.5P (Can add up to 4P when observed in a new family)
>4 patients in the family homozygous or compound heterozygous in *Parkin*? (two unaffected family members with ≤1 variant in *Parkin* considered equivalent to one patient)
4P (Can add up to 4P when observed in a new family)
Unaffected family members homozygous or compound heterozygous with a variant as *pathogenic, likely pathogenic* or *uncertain significance* in *Parkin* at age>50
2B (Can add when observed in a new family)

**Supplementary Figure 1. Flowchart for scoring clinical evidence.** The process of scoring clinical evidence is shown in a root-decision tree flowchart adapted from (1). Different lines of evidence were evaluated to assign pathogenic points (P in pink) and benign points (B in green). **(A)** Minor allele frequency (MAF) from the ExAC population database was used to assign benign and pathogenic points. The cutoffs for each range were established by quantitatively analyzing the frequency spectrum of 1508 known pathogenic variants from 79 disease genes (1). If the variant was absent in ExAC, other public databases were searched. The maximum MAF in these databases was scored accordingly. As indicated in Sherloc, a variant that was found as homozygous in two or more than two individuals in ExAC was attributed 2 benign points. The population cohort in ExAC was not guaranteed to exclude PD patients. However, through multiplying the size of the ExAC cohort (around 60,000 unrelated individuals), by the prevalence of PD in population (~0.3%) (2), the percentage of familial cases (~10%) (3), and the percentage of familial cases caused by *Parkin* (no more than 10%) (3), we estimate a maximum of two patients in ExAC whose PD is caused by recessive pathogenic *Parkin* variants. If two individuals were found in ExAC carrying a homozygous missense variant in *Parkin*, it is very likely that at least one of them did not have PD. According to ACMG-AMP guidelines, a variant observed as homozygous in a healthy adult individual for a recessive (homozygous) disorder, with full penetrance expected at an early age, is likely to be benign and thus 2 benign points were attributed (4). **(B)** Case reports from disease databases reporting families or individuals with PD carrying *Parkin* missense variants were examined. Patients carrying only one variant (heterozygotes) in *Parkin* or more than two variants in *Parkin* or other genes were excluded. For families or unrelated patients homozygous or compound heterozygous for *Parkin* variants, the allele counts of the variants in ExAC were examined. Only when no more than 8 allele counts were reported, could two pathogenic points (homozygote) or 1.5 pathogenic points (compound heterozygous) be attributed for each family or unrelated patient. **(C)** Segregation analysis was performed using family reports with multiple informative members from disease databases. Observation of multiple affected PD patients who were homozygous or compound heterozygous with *Parkin* variants within a family were assigned pathogenic points as shown. Unaffected family members carrying one or no variant in *Parkin* were also scored as shown. Up to four points could be added when segregation was observed within a new family. Benign points were attributed for unaffected family members homozygous or compound heterozygous for *Parkin* variants. Points could also be added when this observation was reported in a new family.

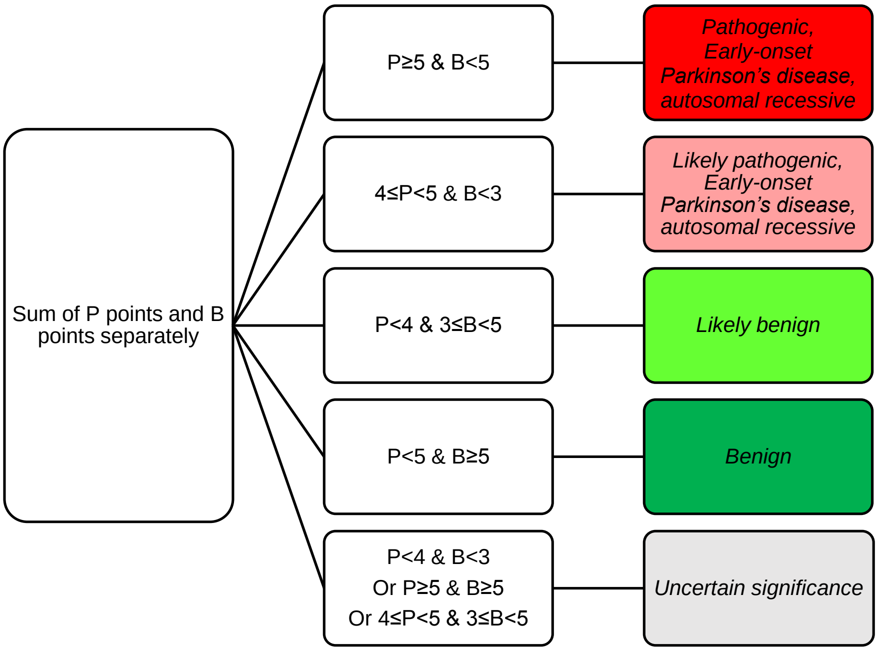


**Supplementary Figure 2. Flowchart for classification of variants.** The process of classification of the pathogenicity of the variants was shown in a root-decision tree flowchart adapted from (1). The benign points and pathogenic points from all evidence were summed separately and compared to the preset threshold. Three benign points was used as the threshold for *likely benign*, and five benign points for *benign*. Four pathogenic points was used as the threshold for *likely pathogenic*, and five for *pathogenic*. When the points exceeded neither the benign nor the pathogenic threshold, the variant was assigned as of *uncertain significance* with insufficient evidence. When the benign and pathogenic scores each exceeded their respective threshold, it suggested the criteria for benign or pathogenic was conflicting. The variant was also assigned as of *uncertain significance*, with indication of low-penetrance variants, genetic or environmental modifiers, or other ambiguity within the Mendelian framework (1).

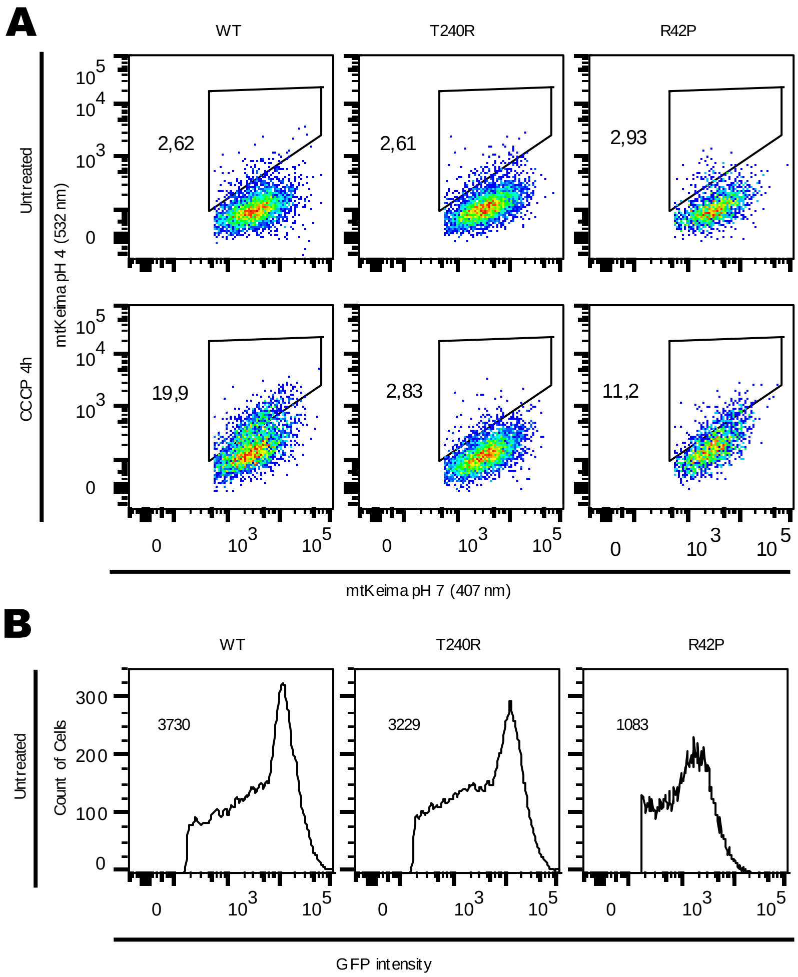


**Supplementary Figure 3. Mitophagy and steady-state Parkin protein levels were examined using a FACS-based analysis of mitochondrially-targeted mKeima (mtKeima) and GFP intensity. (A)** Representative FACS data of mt-mKeima expressing GFP fused WT Parkin or Parkin variants, untreated or treated with CCCP for 4 h. Mitophagy was calculated as the percentage of cells in the gate of enhanced mtKeima fluorescent signal at pH4 versus pH7. **(B)** Representative histogram of GFP signal in cells expressing GFP fused WT or Parkin variants (without CCCP treatment). Geometric mean of GFP signal was calculated from the histogram.

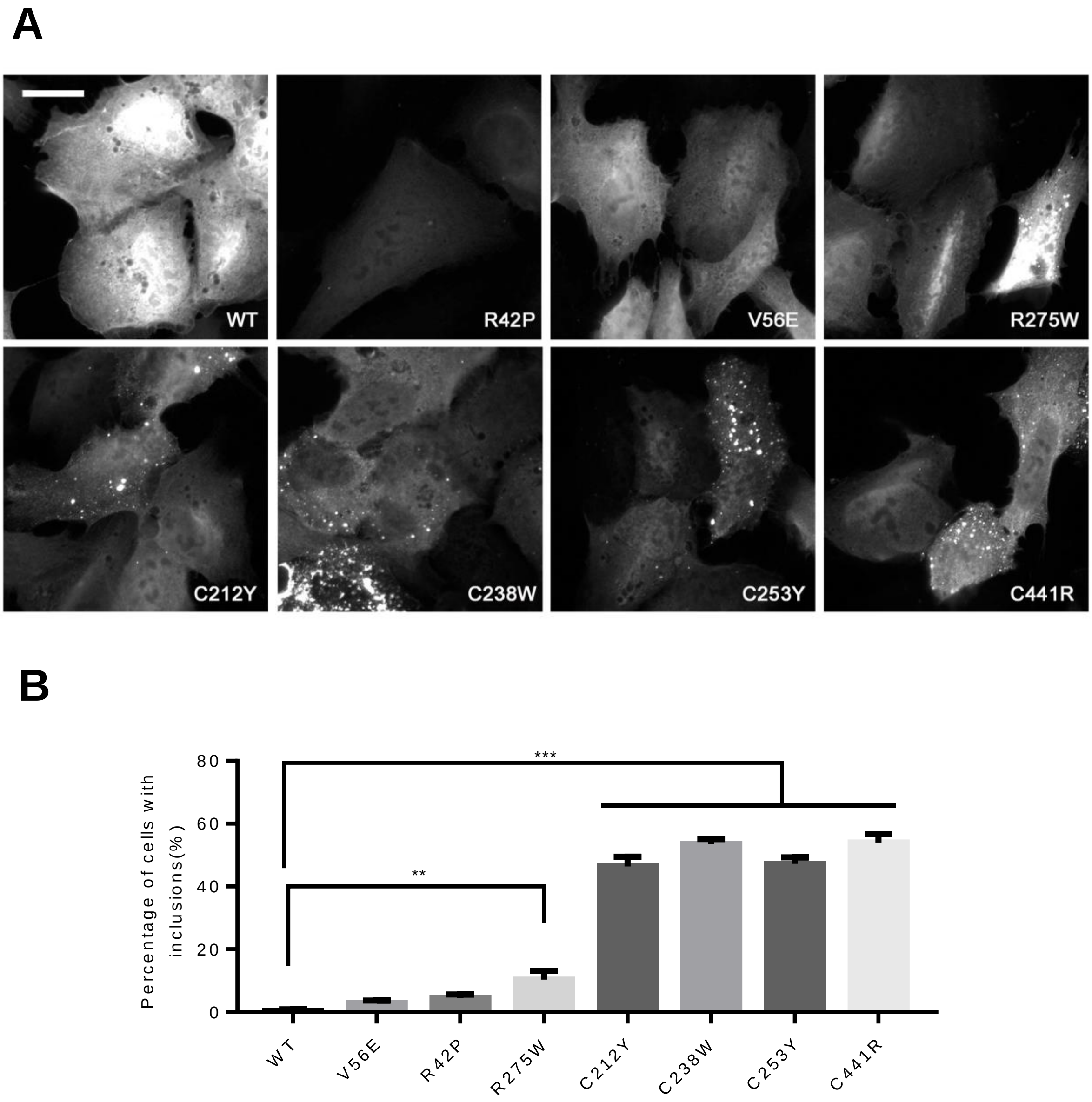


**Supplementary Figure 4. Parkin variants in the R0RBR form inclusions. (A)** Representative image of GFP distribution in U2OS cells expressing GFP-Parkin WT or the indicated Parkin variants. Scale bars, 20 μm. **(B)** Quantification of percentage of cells with more than 2 inclusions. **P<0.01, ***P<0.001. N=5 groups with 50 cells per group. One-way ANOVA with Dunnett's post-hoc test comparing each variant with WT.

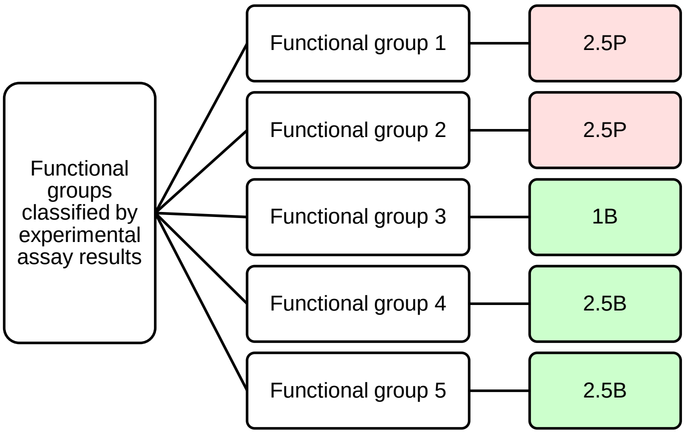


**Supplementary Figure 5. Flowchart for scoring functional evidence.** The process of scoring the functional evidence was shown in a root-decision tree flowchart adapted from (1). Points were assigned to the variants according to their classification to the functional groups from Figure 4A.

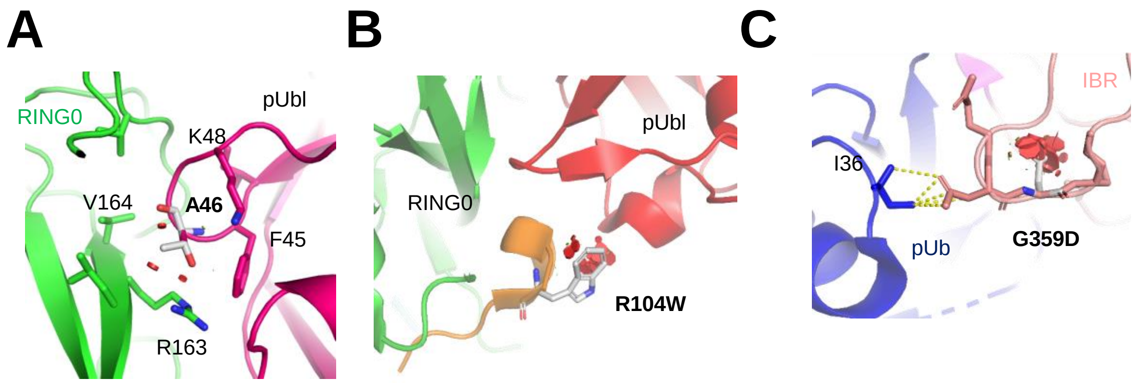


**Supplementary Figure 6. Structural simulation of *Parkin* variants. (A)** Close-up view of pUbl-RING0 interface in structure of fly pParkin bound to pUb (PDB 6DJX). The A46T variant in the pUbl would disrupt the interaction with RING0 by introducing clashes with Val164 and Arg163. **(B)** Close-up view of the ACT-pUbl-RING0 interface in the structure of human pParkin bound to pUb (PDB 6GLC). The R104W variant introduced major clashes that would disrupt the interaction of pUbl with RING0. **(C)** Close-up view of IBR-pUb interface in structure of fly pParkin bound to phospho-ubiquitin (PDB 6DJX). The G359D variant would introduce major clashes to the loop in the IBR domain that interacts with pUb.

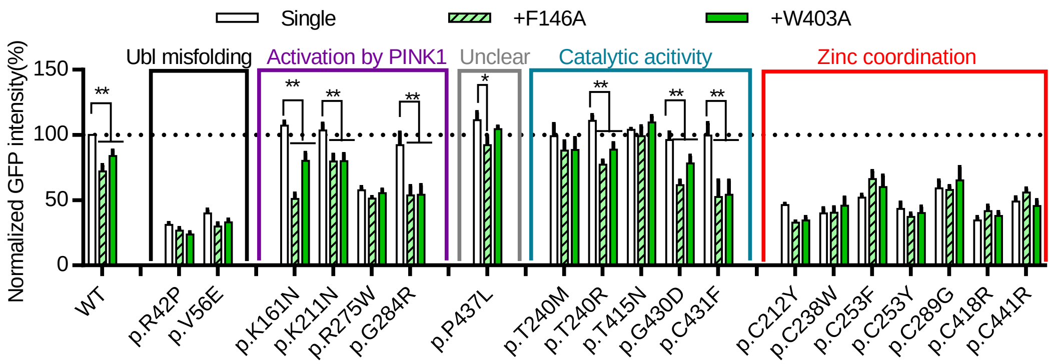


**Supplementary Figure 7. Effects of designer activating mutations on WT and variant Parkin levels.** GFP intensity was quantified by FACS in untreated cells expressing GFP fused with WT Parkin, *pathogenic* variants, W403A, F146A or variants *in cis* with W403A or F146A. The GFP intensity for each Parkin missense variant was normalized to that for WT Parkin in each replicate. *P<0.05, **P<0.01, in two-way ANOVA with Dunnett's post-hoc test comparing the function of each variant with the variant *in cis* with W403A or F146A. N=3-7.

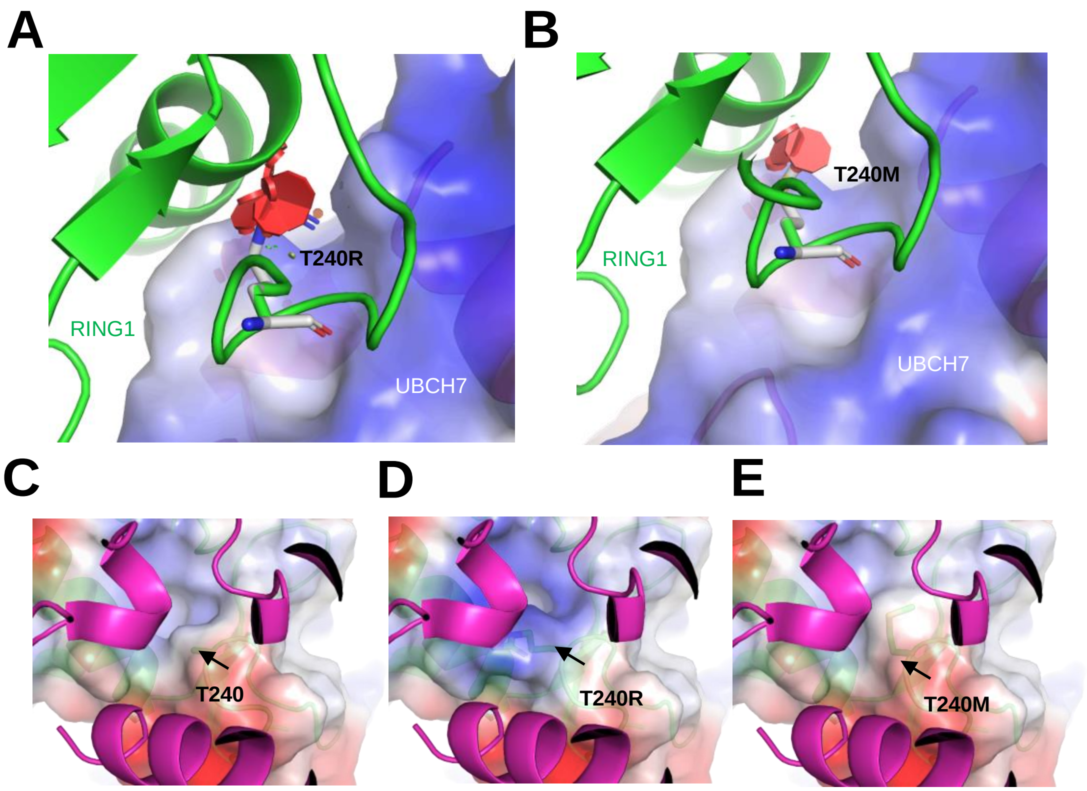


**Supplementary Figure 8. Structural simulations reveal the effects of the different T240 variants on E2 binding. (A-B)** Close-up view of RING1:UBCH7 interface in the structure of fly pParkin bound to pUb and UBCH7 (PDB 6DJW). The electrostatic surface of UBCH7 at the interface was shown (Blue, positive charge). Mutation of T240 (A275) to **(A)** arginine (stick) or **(B)** methionine (stick) introduced major clashes (red disks) at the interface and thus is predicted to weaken the interaction with E2. **(C-E)** Close-up view of RING1:UBCH7 interface in the structure of fly pParkin bound to pUb and UBCH7 (PDB 6DJW). The electrostatic surface of RING1 domain at the interface is shown (Blue, positive charge). Mutation of T240 (A275) to arginine (stick) creates a positive charge at the interface (indicated with the arrow), increasing the electrostatic repulsion of UBCH7. **(C)** T240 (A275). **(D)** T240R (A275R). **(E)** T240M (A275R).

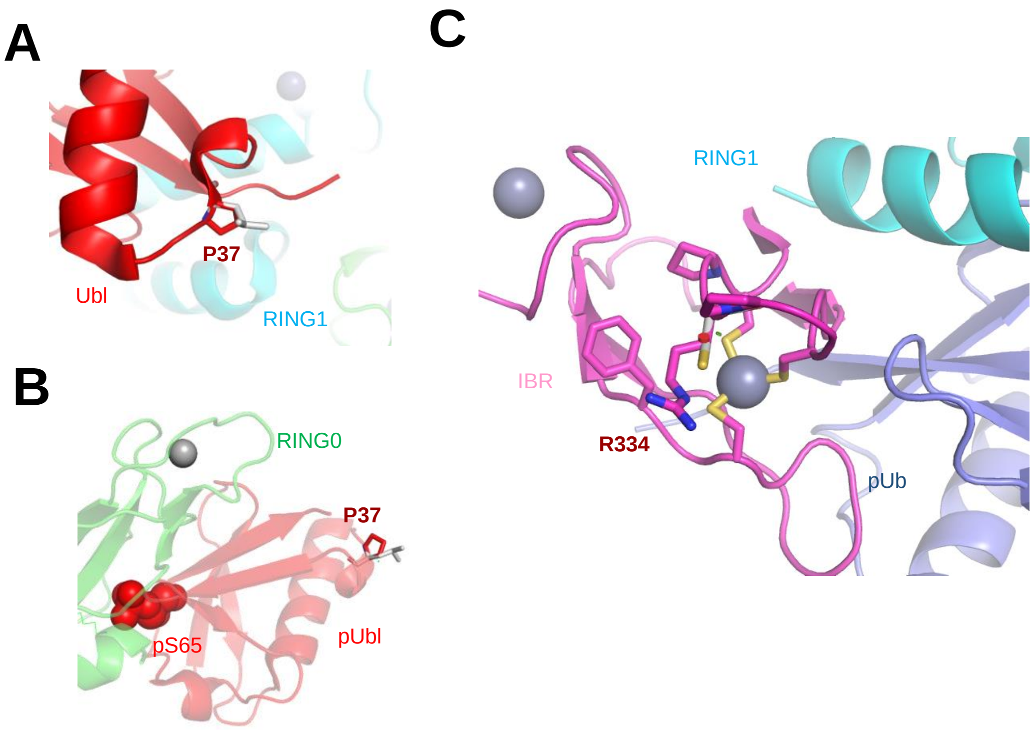


**Supplementary Figure 9. Structural basis for the effects of naturally occurring hyperactive Parkin variants. (A)** Close-up view of the P37L variant site (PDB 5N2W). A leucine at this position (white sticks) would not introduce a steric clash and would not affect interaction of the Ubl with RING1. **(B)** Close-up view of the P37L variant site (PDB 6GLC). A leucine at this position (white sticks) would not introduce a clash and would not affect interaction of the pUbl with RING0. **(C)** Close-up view of R334C variant site (PDB 5N2W). A cysteine at this site (white stick) could coordinate the nearby zinc, thus further stabilizing the interaction with pUb.


1. Nykamp K, Anderson M, Powers M, Garcia J, Herrera B, Ho Y-Y, et al. Sherloc: a comprehensive refinement of the ACMG–AMP variant classification criteria. *Genet Med.* 2017;19:1105.
2. Poewe W, Seppi K, Tanner CM, Halliday GM, Brundin P, Volkmann J, et al. Parkinson disease. *Nature Reviews Disease Primers.* 2017;3:17013.
3. Klein C, and Westenberger A. Genetics of Parkinson's Disease. *Cold Spring Harbor Perspectives in Medicine.* 2012;2:a008888.
4. Richards S, Aziz N, Bale S, Bick D, Das S, Gastier-Foster J, et al. Standards and guidelines for the interpretation of sequence variants: a joint consensus recommendation of the American College of Medical Genetics and Genomics and the Association for Molecular Pathology. *Genet Med.* 2015;17:405-24.
